# Thermodynamic destabilization informs pathogenicity assessment of a variant of uncertain significance in cardiac myosin binding protein C

**DOI:** 10.1101/789081

**Authors:** Maria Rosaria Pricolo, Elías Herrero-Galán, Cristina Mazzaccara, Maria Angela Losi, Jorge Alegre-Cebollada, Giulia Frisso

## Abstract

In the era of Next Generation Sequencing (NGS), genetic testing for inherited disorders identifies an ever-increasing number of variants whose pathogenicity remains unclear. These variants of uncertain significance (VUS) limit the reach of genetic testing in clinical practice. The VUS for Hypertrophic Cardiomyopathy (HCM), the most common familial heart disease, constitute over 60% of entries for missense variants shown in ClinVar database. We have studied a novel VUS (c.1809T>G-p.I603M) in the most frequently mutated gene in HCM, *MYBPC3*, which codes for cardiac myosin-binding protein C (cMyBPC). Our determinations of pathogenicity integrate bioinformatics evaluation and functional studies of RNA splicing and protein thermodynamic stability. In silico prediction and mRNA analysis indicated no alteration of RNA splicing induced by the variant. At the protein level, the p.I603M mutation maps to the C4 domain of cMyBPC. Although the mutation does not perturb much the overall structure of the C4 domain, the stability of C4 I603M is severely compromised as detected by circular dichroism and differential scanning calorimetry experiments. Taking into account the highly destabilizing effect of the mutation in the structure of C4, we propose reclassification of variant p.I603M as likely pathogenic. Looking into the future, the workflow described here can be used to refine the assignment of pathogenicity of variants of uncertain significance in *MYBPC3*.

## 1. Introduction

Hypertrophic cardiomyopathy (HCM) constitutes the most common inherited disease of the myocardium with an estimated prevalence ranging from 1:500 to 1:200 ^1,2^. Although the clinical manifestations of HCM are highly variable, the disease is characterized by left ventricular hypertrophy (LVH) in the absence of triggers, such as hypertension, and in some cases sudden cardiac death (SCD) may be the first manifestation of the disease ^3–5^. Importantly, HCM has a very strong genetic component, and up to 60% of patients who meet the diagnostic criteria have pathogenic mutation in genes coding for sarcomere proteins ^6–8^. The genetic testing for HCM is key to identify patients at high risk before the occurrence of clinical manifestations. Thus, the clinical role of genetic testing largely centers on family screening to facilitate presymptomatic diagnosis of family members, clinical surveillance and reproductive advice ^9–12^. Due to the increased complexity of analysis and interpretation of genetic tests, general guidelines for the interpretation of variants have been published by the American College of Medical Genetics and Genomics (ACMG) and the Association for Molecular Pathology (AMP) ^13,14^. By applying the proposed score, variants can be classified into five main groups including pathogenic, likely pathogenic, likely benign, benign, and uncertain significance variants (VUS). VUS hamper clinical interpretation and risk-assessment limiting counselling and treatment of carriers ^15–17^. The numbers of mutations classified as VUS has increased in the last years ^18^, reaching 66.5% of entries in ClinVar database for HCM missense mutations. To improve care of many HCM patients and their families, there is a pressing need to develop methods that can help reclassification of VUS into actionable categories. According to ACMG, in the absence of enough genetic support, functional studies are the most important criterion to establish pathogenicity of putative disease-causing mutations ^14^.

Mutations in the *MYBPC3* gene, encoding cardiac myosin-binding protein C (cMyBPC), represent 40–50% of all HCM mutations, making it the most frequently mutated gene in this disease ^19–21^. *MYBPC3* is also the HCM gene with the highest number of missense VUS in ClinVar. *MYBPC3* pathogenic missense mutations can lead to stable mutant cMyBPCs that are, at least in part, incorporated into the sarcomere and could act as poison polypeptides by alteration of the structure and/or function of the sarcomere ^21^. Missense mutations located in the central domains (C3-C7) of the protein, and clinically causative of HCM, have been identified to interfere with the structure and the stability of the domains ^22,23^. Other pathogenic missense mutations of cMyBPC, are known to cause RNA splicing disruptions ^24,25^. Interestingly, both protein destabilization and alterations of splicing can lead to less cMyBPC levels, a well-known driver of HCM ^26–28^.

During genetic screening of HCM patients, we found a new putative missense mutation in the *MYBPC3* gene (c.1809T>G-p.I603M), which targets the C4 central domain of cMyBPC. Here, we describe how available population and computational data result in the classification of this variant as VUS according to ACMG criteria. We then demonstrate that this variant induces extensive protein destabilization, which leads to reclassification of c.1809T>G-p.I603M as likely pathogenic.

## 2. Methods

### 2.1 Molecular genetics of HCM patients

Human samples to perform genetic analysis were obtained following informed consent of patients according to the Declaration of Helsinki. Genomic DNA was isolated from peripheral whole blood as previously described ^29^. All coding exons, and 5′ and 3′ UTRs of genes involved in HCM were amplified by PCR and analysed by automatic sequencing using previously reported protocols ^9^.

### 2.2 RNA Splicing predictions and analysis

Alamut software (Alamut® Visual, Interactive Biosoftware) was used for *in silico* prediction of splice-affecting nucleotide variant ^30,31^. Genomic sequences (WT and mutant) were processed using five splicing prediction tools (SpliceSiteFinder-like, MaxEntScan, Neural Network Splice, GeneSplicer, and Human Splicing Finder). RNA splicing was experimentally examined using mRNA from peripheral blood. Total RNA was extracted from lymphocyte cells of patients’ peripheral blood using Trizol Reagent (Thermo Fischer Scientific, Waltham, MA, USA). RNA retro-transcription was performed by SuperScript VILO (Life Technologies), starting from 1μg of total RNA and using random primers. The cDNA obtained was amplified by PCR using specific consecutive-exon-spanning primers: **MYBPC3_ex15_Fw** 5’-CAAGCGTACCCTGACCATCA-’3 and **MYBPC3_ex20-21_Rv** 5’-GGATCTTGGGAGGTTCCTGC-’3 oligonucleotides which anneal with exon 15 and the region encompassing exons 20 and 21 of the *MYBPC3* mRNA, respectively. The same primers were used for sequencing the PCR fragment. Sequences were analysed with CodonCode Aligner software (CodonCode Corporation, Dedham, MA, USA).

### 2.3 Bioinformatics predictions of pathogenicity at the protein level

Missense mutation at the protein level was evaluated with three independent bioinformatics tools that predict possible impact of amino acid substitutions on the structure and function of the protein (PolyPhen-2, SIFT, Provean). **PolyPhen-2** (Polymorphism Phenotyping v2) evaluates the impact of amino acid allelic variants via analysis of multiple sequence alignments and protein 3D-structures ^32^. **SIFT** (Sorting Intolerant From Tolerant) presumes that important amino acids will be conserved in the protein family, so changes at well-conserved positions tend to be predicted as deleterious ^33^. **PROVEAN** (Protein Variation Effect Analyzer) predicts whether an amino acid substitution has an impact on the biological function of a protein using pairwise sequence alignment scores ^34^.

### 2.4 Homology modelling

For modelling of cMyBPC’s C4 domain, the protein sequence Q14896 from the UniProt database was used (aa.544-aa.633). The structure was modelled using I-TASSER tool (Iterative Threading ASSEmbly Refinement) ^35^, indicating immunoglobulin domain four of slow-MyBPC (2YUZ PDB) as the best template. PyMol software was used for molecular representation ^36^.

### 2.5 Protein expression and purification for biophysical characterization

The recombinant WT and mutant I603M C4 domains (Table 1) were engineered and purified for thermodynamic analysis. With that aim, the cDNA fragment including exons 18 and 19 of MYBPC3, was amplified from cardiac RNA with **cMyBPC_C4_Fw** and **cMyBPC _C4 _Rv** oligonucleotides (Table 2) and cloned in a custom-modified pQE80L (Qiagen) using BamHI and BglII restriction sites. I603M mutant cDNA was generated by PCR site-directed mutagenesis. Final constructs were verified by Sanger sequencing. Proteins were expressed in BLR (DE3) *E. coli* strain and purified from the soluble fraction using nickel-nitrilotriacetic acid (Ni-NTA) chromatography (Qiagen) with a column volume of 3 mL and pre-equilibrated with phosphate buffer (50 mM sodium phosphate pH 7 and 300 mM NaCl) supplemented with 10 mM DTT. Elution was performed in two steps with increasing imidazole concentration (from 20mM to 250mM). Further purification was achieved by size-exclusion chromatography in an AKTA Pure 25L system using a Superdex 200 Increase 10/300 GL column (GE Healthcare). The proteins were eluted in phosphate buffer buffer (20 mM sodium phosphate pH 6.5, 50 mM NaCl), which was used both in circular dichroism (CD) and differential scanning calorimetry (DSC) experiments. All domains were at least 90–95% pure as estimated by SDS-PAGE analysis.

**Table 1:** Properties of recombinant C4 WT and I603M. Sequences corresponding to C4 are indicated in bold whereas extra amino acids resulting from cloning are shown in regular type. Predicted extinction coefficients were obtained from ProtParam tool^59^.

| Protein | Sequence | Number of amino acids | Extinction coefficient (cm<sup>2</sup> x mg<sup>-1</sup>) |
| --- | --- | --- | --- |
| C4 WT | MRGSHHHHHHGS**KLEVYQSIADLMVG**<br>**AKDQAVFKCEVSDENVRGVWLKNGKE**<br>**LVPDSRIKVSHIGRVHKL**TIDDVTPADE<br>ADYSFVPEGFACNLSAKLHFMERS | 104 | 0.736 |
| C4 I603M | MRGSHHHHHHGS**KLEVYQSIADLMVG**<br>**AKDQAVFKCEVSDENVRGVWLKNGKE**<br>**LVPDSRIKVSHIGRVHKL**TMDDVTPAD<br>EADYSFVPEGFACNLSAKLHFMERS | 104 | 0.724 |

**Table 2:**
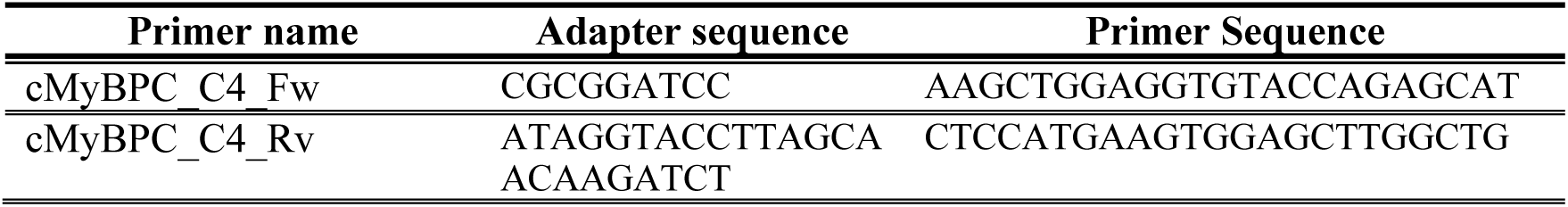
Primers used to clone C4 (5’-3’)

### 2.6 Circular Dichroism (CD)

CD spectra of domain C4 were collected with a Jasco-810 spectropolarimeter. The temperature was controlled using a Peltier thermoelectric system. Purified proteins in phosphate were loaded into a 0.1cm or 1cm path-length quartz cuvettes for data collection in the far-UV (195-250 nm) or the near-UV (250-350nm), respectively. Protein concentration was 0.3 mg/mL (far-UV) and 1 mg/mL (near-UV). Spectra were collected for the protein samples and buffer control at 25°C and 85°C with four accumulations of data. The buffer baseline spectrum was subtracted from each protein spectrum at each corresponding temperature to correct for the background signal. Thermal denaturation analyses were carried out collecting the variations of ellipticity at 230 nm as a function of temperature (25°C-85°C, at a rate of 30°C/h). Protein refolding was analysed with a temperature ramp-down (85°C-25°C) at the same speed. We plotted the ellipticity versus temperature and the data points were fit with a Boltzmann sigmoidal curve using Igor Pro software to obtain midpoint unfolding temperatures (T_m_). The changes in CD as a function of temperature can be used to determine the van’t Hoff enthalpy (ΔH_v_) of unfolding considering ΔCp = 0 ^37^.

### 2.7 Differential scanning calorimetry (DSC)

Calorimetric measurements were performed using a Microcal VP-DSC differential scanning calorimeter with 0.5 mL cells. Experiments were done by increasing temperature from 25°C to 85°C at a rate of 30°C/h, using 0.085 mM (1 mg/mL) protein concentration in phosphate buffer. The reversibility of the thermal transitions was assessed by reheating of the sample immediately after cooling from the previous scan, using the same rate of temperature change. The calorimetric traces were corrected for the instrumental background by subtracting a scan with buffer in both cells. The temperature dependence of the excess heat capacity was analysed using Origin software (MicroCal, Northampton, MA). The thermal stability of the proteins was described by its T_m_, and the calorimetric enthalpy (ΔH_cal_), which was calculated as the area under the excess heat capacity function. The van’t Hoff enthalpy (ΔH_v_) associated with DSC thermograms was determined by a two-state fit of the thermograms ^38,39^. The Gibbs free energy change (ΔG) was then calculated at any temperature using three experimental parameters: T_m_, enthalpy change at T_M_ (ΔH(T_m_)) and heat capacity change at T_m_ (ΔC_p_). The ΔG was derived for C4 WT and C4 I603M considering ΔC_p_ = 0.

## 3. Results

### 3.1 c.1809T>G (p.I603M) is classified as VUS according to available population and bioinformatics predictions

The genetic screening of an HCM patient highlighted a missense variant in MYBPC3 (c.1809T>G-p.I603M) that was not reported before as associated with HCM. The variant occurs in exon 19 of the gene, whereas at the protein level the Ile603 maps to the central domain C4 of cMyBPC. The p.I603M substitution was identified in a proband with family history of HCM (Table 3). Although segregation analysis was possible, the family tree was not informative (*Figure 1, A*). The genetic screening of other family members evidenced the presence of the c.1809T>G-p.I603M variant in two affected subjects in heterozygosis with another MYBPC3 mutation, p.T33RfsX15. One subject (III.5 in *Figure 1, A*) is clinically affected but did not carry the p.I603M mutation, although hypertrophy in this individual can be secondary to coarctation of the aorta. Two young subjects carry the c.1809T>G-p.I603M variant and show so far no clinical evidence of HCM (IV.1 and IV.2 in *Figure 1, A;* Table 3).

**Table 3:** Clinical data.

| PATIENT | AGE AT ONSET | SYNCOPE | CONDUCTION DEFECTS | ARRHYTHMIAS | LVEDD (mm) | LVESD (mm) | EF (%) | MWT (mm) |
| --- | --- | --- | --- | --- | --- | --- | --- | --- |
| III.1 | 39 | - | LAH | - | 40 | 23 | 70 | 26 |
| III.2 | asymptomatic | - | - | - | 45 | 29 | 56 | 10 |
| III.3* | 17 | - | RBBB | NSVT | 48 | 27 | 58 | 19 |
| III.5 | 21 | - | - | - | NR | NR | 65 | 19 |
| IV.1 | asymptomatic | - | - | - | 42 | 21 | 57 | 9 |
| IV.2 | asymptomatic | - | - | - | 44 | 23 | 65 | 7 |

\*proband; LVEDD: left ventricular end diastolic diameter; LVESD: left ventricular end systolic diameter; EF: ejection fraction; MWT: maximal wall thickness; RBBB: right bundle branch block; LAH: left anterior hemiblocks; NSVT: nonsustained ventricular tachycardia; NR: not reported

**Figure 1:**
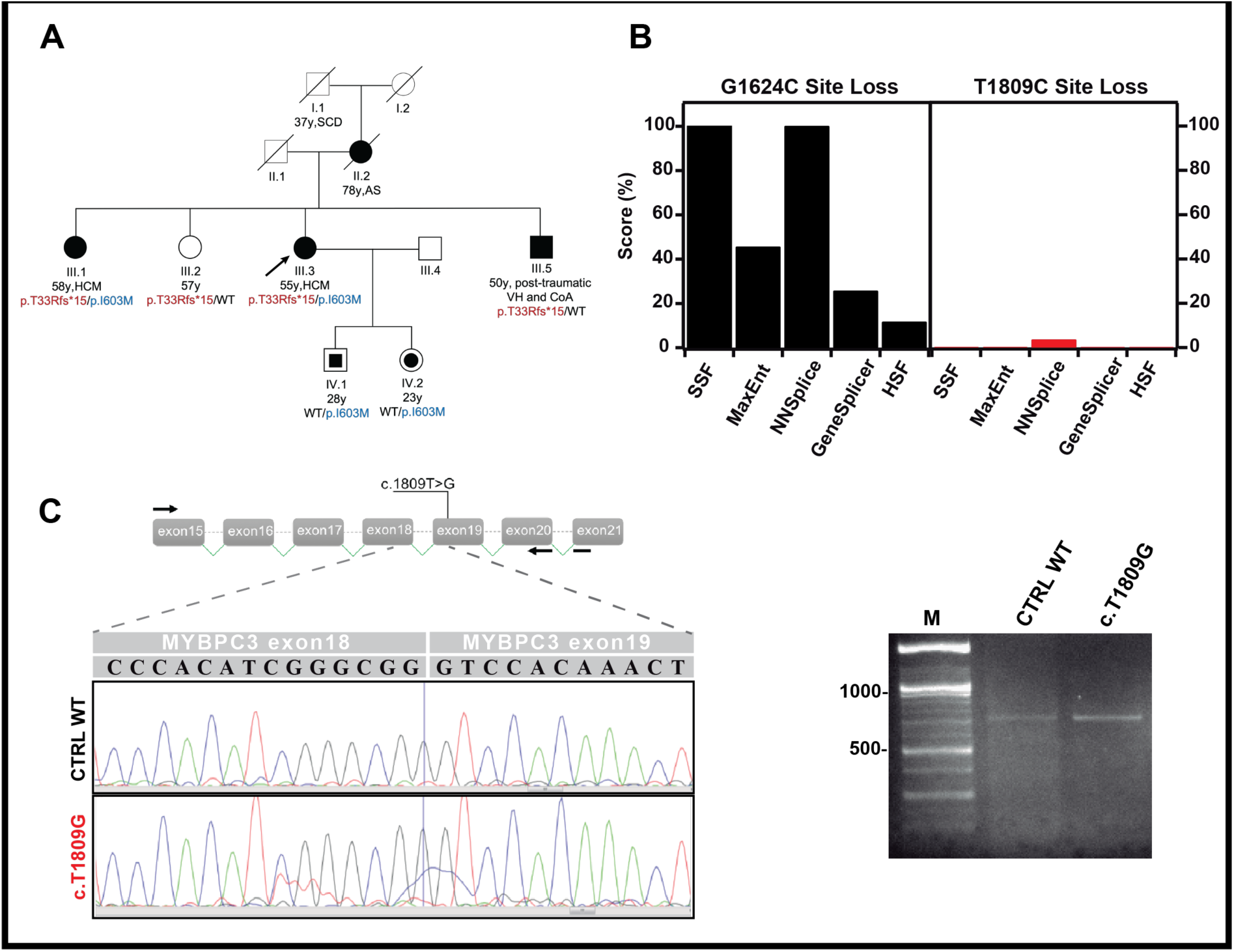
Study of the potential effect of the MYBPC3-c.1809T>G (p.I603M) mutation on RNA splicing. **(A)** Pedigree of four generations of the HCM family with the p.I603M variant. *Open symbols* represent subjects with a negative phenotype. *Filled symbols* represent clinically affected subjects. *Circles or square with solid centres* indicate unaffected mutation carriers. Black arrow indicates proband. Relevant clinical features are indicated: SCD, Sudden Cardiac Death; AS, Aortic Stenosis; HCM, Hypertrophic cardiomyopathy; VH, Ventricular Hypertrophy; CoA, coarctation of the Aorta. y:years. **(B)** Prediction of RNA splicing alterations using Alamut software. The histograms represent the score given by the different prediction algorithms for a splicing site loss calculated between WT and the corresponding mutation. The evaluations were carried out with five algorithms of splicing prediction: SSF (SpliceSiteFinder-like), MaxEnt, NNSPLICE (Neural Network Splice), GeneSplicer and HSF (Human Splicing Finder). Predictions for the c.1624G>C (E542Q) mutation (*left*). Prediction for c.1809 T>C (p.I603M) variant (*right*). For mutation c.1624G>C (E542Q), the loss of the canonical splicing site is detected by all the algorithms. For variant c.1809 T>C (p.I603M), the prediction suggests no RNA splicing alteration. (**C**) *Top left*: Experimental study of RNA splicing by RT-PCR analysis of mRNA isolated from peripheral blood. The MYBPC3 fragment analyzed is represented. Black arrows indicate forward and reverse PCR oligonucleotides. *Bottom left*: Electropherograms obtained from cDNA sequencing of PCR fragments. CTRL WT corresponds to the analysis of mRNA obtained from a healthy, non-carrier subject. The electropherograms indicate the correct joining of exons 18-19 in both samples. (*Right*) Electrophoresis of RT-PCR analyses of mRNA. Lane 1: Marker XIV (Roche Diagnostics); Lane 2: Non-carrier control; Lane 3: c.1809T>G mutation.

We analysed the very strong ACMG criteria of pathogenicity corresponding to population frequency and predictors of pathogenicity. We first evaluated Minor Allele Frequency (MAF) in ExAC and GnomAD databases ^40,41^. The MAF of allele c.1809G is 0.00004267 and 0.0000142 in ExAC and GnomAD, respectively. These MAF values indicate that c.1809T>G is a rare variant in the population, compatible with HCM ^42^. Evaluation using Polyphen, SIFT and PROVEAN tools showed strong evidence for the deleterious effect of the p.I603M variant (Table 4). Indeed, in comparison with the non-pathogenic variant p.G278E ^43,44^ in MYBPC3, p.I603M showed a higher score of pathogenicity. According to ACMG criteria, in particular with respect to the low MAF and the results from the bioinformatics predictors, the variant c.1809T>G (p.I603M) is classified as VUS. Hence, we experimentally determined if the mutation can affect the bioavailability of cMyBP-C at the level of alterations of RNA processing or protein stability, as reported for other mutants ^21–25,45,46^.

**Table 4:** bioinformatic predictions of p.I603M deleterious effect.

| **cMyBPC**<br>**variant** | **PolyPhen-2**<br>($> 0.6$) | **SIFT**<br>($< 0.05$) | **PROVEAN**<br>($< -2.5$) |
| --- | --- | --- | --- |
| p.I603M<br>(c.1809 T>G) | 1.0 | 0.001 | -2.63 |
| p.G278E<br>(c.833 G>A)* | 0.198 | 0.084 | -1.79 |

Numbers in parentheses indicate the threshold of each predictor to assign a deleterious effect \* dbSNP code: rs147315081

### 3.2 c.1809T>G (p.I603M) does not induce alterations in RNA splicing

The removal of introns from pre-mRNA and the joining of exons are critical aspects of gene expression. Introns are removed from primary transcripts by the process of RNA splicing, which links together the flanking exons to generate mRNA ^47^. We investigated potential RNA splicing alterations induced by variant c.1809T>G-p.I603M. Bioinformatics predictions did not suggest loss of natural splice sites. Indeed, all five algorithms did not show any probability of canonical splice site loss compared to a well-established pathogenic mutation c.1624G>C (p.E542Q) of *MYBPC3* ^27,48^ (*Figure 1, B*). Similar results were obtained for the prediction of activation of cryptic splicing sites (data not shown). To verify predictions, we amplified the fragment spanning from exon 15 to exon 21 using mRNA isolated from peripheral blood. The gel electrophoresis showed an identical migration pattern of both mutant carrier and a healthy control (*Figure 1, C right*). Sanger sequencing confirmed the correct splicing in both cases (*Figure 1, C left*).

### 3.3 C4 WT and C4 I603M share a similar β-sheet rich structure

The p.I603M is a missense variant that affects a highly conserved residue in the central domain C4 of cMyBPC (*Figure 2, A, D)*. Since the structure of domain C4 is not known, to predict how the protein structure might be affected by the p.I603M mutation, the homology modelling of the domain was carried out using I-TASSER. The resulting model showed that the C4 domain adopts an Ig-like folding (*Figure 2, B*) ^49^. The predicted structure of C4 I603M is very similar to WT. The RMSD of both predictions with respect to the template is 1.9±1.6Å for WT and 2.0±1.6Å for p.I603M. The RMSD of WT and p.I603M homology models is 0.2634Å.

**Figure 2:**
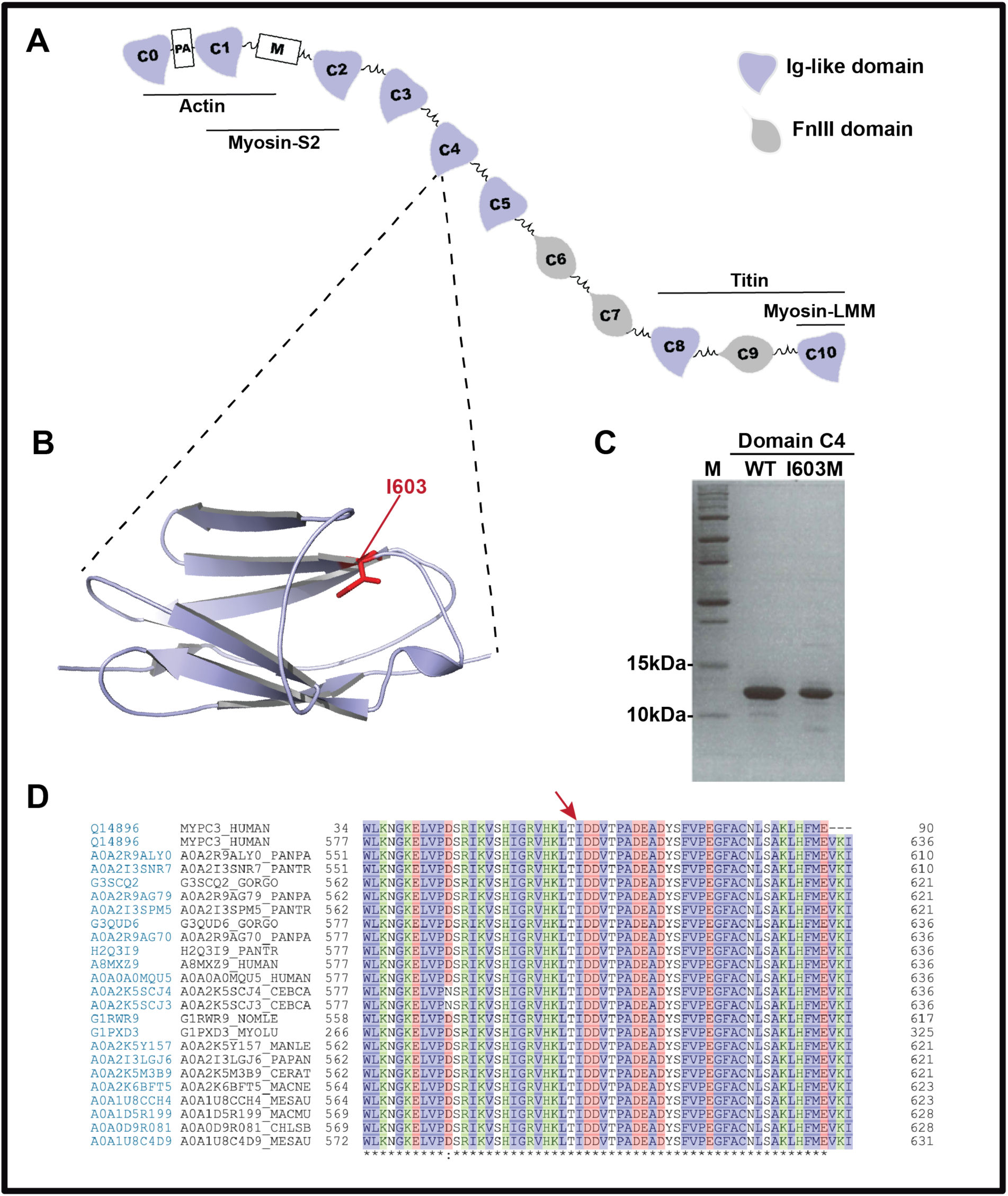
MYBPC3-c.1809T>G (p.I603M) mutation at the protein level. (**A**) Schematic representation of cardiac Myosin binding protein C (cMyBPC) and its sarcomeric interactor. cMyBPC is a multimodular protein composed by a series of immunoglobulin-like (*violet*) and fibronectin-type III (*gray*) domains. PA correspond to Pro/Ala rich region between C0 and C1. A 105-residue linker between C1 and C2 is called M-motif (M). The NH2-terminus of cMyBPC interacts with Actin (C0-M) and the myosin-S2 domain (C1-M-C2). The COOH-terminus of cMyBPC interacts with titin (C8-C10) and light meromyosin (C10). (**B**) Homology modelling prediction of cMyBPC C4 domain. Graphic representation of the 3D structure of C4 WT domain, modelled with I-tasser using slow-MyBPC as a template (2YUZ PDB structure). The position of the isoleucine replaced in HCM patients is shown in red. The models of C4 domain have been prepared with PyMOL. (**C**) SDS-PAGE analysis of purified recombinant C4 domain expressed in *E. coli* BLR21 (DE3). (**D**) Sequence alignment of human cMyBPC C4 domain with other species. Data from UNIPROT were aligned by ClustalW. Regions between amino acids 34-87 of C4 domain are shown. The red arrow indicates position of Ile 603.

Both isoleucine and methione share similar physico-chemical properties, suggesting no effect on protein structure by mutation p.I603M. Still, it is interesting to observe that Ile603 is buried in the core of the WT domain (*Figure 2, B*), so a mutation affecting this residue may not be easily accommodated leading to destabilization. To study this scenario, recombinant WT and I603M C4 domains were produced (*Figure 2, C*) and analysed by Circular Dichroism (CD) in the far-UV (200-250 nm, informative about secondary structure) and near-UV (250-350 nm, resulting from tertiary structure) at 25°C (*Figure 3 A, B*). In agreement with the homology modelling prediction, the far-UV spectrum of C4 WT showed a minimum of ellipticity at 215 nm, characteristic of β-sheet structure ^50,51^. C4 WT and I603M spectra showed similar CD spectra, suggesting that the mutation did not have major impact in the fold of the protein at 25°C (*Figure 3, A, B*).

**Figure 3:**
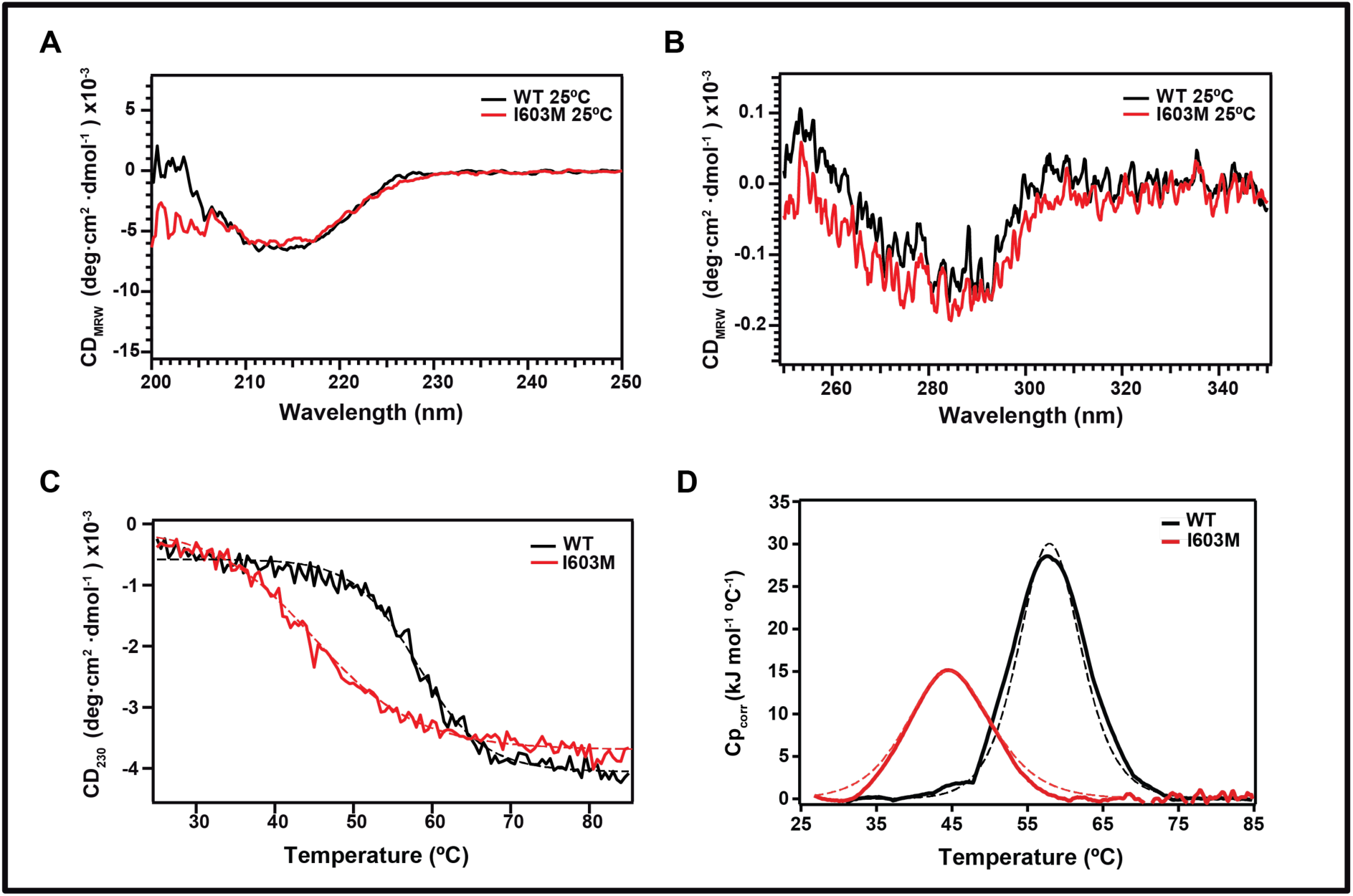
The I603M mutation leads to thermodynamic destabilization of the C4 domain of cMyBPC. (**A**) Far-UV CD spectra of native cMyBPC C4 WT (*black*) and I603M (*red*) monitored at 25°C. (**B**) Near-UV CD spectra of native C4 WT (*black*) and C4 I603M (*red*). (**C**) Thermal denaturation curves obtained for C4 WT (*black*) and I603M (*red*) by tracking CD signal at 230 nm as a function of temperature. The temperature at the midpoint of the transition, T_m_ is obtained by performing a sigmoidal fitting to denaturation curves (*dashed lines*) considering a two-state unfolding process. The domain C4 WT shows a T_m_ of 58°C, 13°C higher than C4 I603M. (**D**) Thermal unfolding of C4 WT and C4 I603M monitored by DSC. The thermograms represent processed experimental data after baseline subtraction. The calorimetric enthalpy was calculated by integration of the area under the thermogram peak. The van’t Hoff enthalpy was determined by fit two a two-state model of unfolding (*dashed lines*).

### 3.4 The mutation p.I603M destabilizes the C4 domain

Thermodynamic protein stability was investigated by monitoring the CD signal at 230nm while increasing temperature from 25°C to 85°C. The unfolding profiles showed a T_m_ of 58°C for C4 WT and 45°C for the I603M mutant (*Figure 3, C; Supplementary Figure 1;* Table 5). A 13°C decrease in T_m_ highly suggests that the I603M mutation induces considerable destabilization of the C4 domain.

**Table 5:** thermal stability parameters calculated by CD and DSC.

|  | **T<sub>m</sub> CD** | **ΔH<sub>v</sub> CD<sub>230</sub>** | **T<sub>m</sub> DSC** | **ΔH<sub>cal</sub> DSC** | **ΔH<sub>v</sub> DSC** |
| --- | --- | --- | --- | --- | --- |
| **C4 WT** | 58 °C | 220 kJ/mol | 58°C | 364 kJ/mol | 330 kJ/mol |
| **C4 I603M** | 45 °C | 130 kJ/mol | 45°C | 196 kJ/mol | 222 kJ/mol |

For further thermodynamic characterization, we undertook refolding CD experiments by ramping the temperature down after the initial heating ramp. The refolding ability of the domains was determined by collecting a final far-UV CD spectrum at 25°C. Most protein refolded into original secondary structure for both WT and I603M (*Supplementary Figure 2*), suggesting that thermal denaturation of both proteins is a reversible process. We used the Gibbs-Helmholtz equation to fit the change of ellipticity at 230 nm as a function of temperature, which enabled estimation of the van’t Hoff enthalpies (ΔH_v_). We obtained that ΔH_v_ of C4 WT is 220 kJ/mol, whereas the value for I603M is 130 kJ/mol for C4 I603M (Table 5). Such difference in enthalpies shows considerable thermodynamic destabilization induced by the I603M mutation.

When the unfolding of a protein is reversible and two-state, the thermodynamic parameters evaluated by CD are almost identical to the ones estimated by calorimetric methods ^37^. However, to confirm thermodynamic destabilization, differential scanning calorimetry (DSC) experiments were carried out in the same conditions as CD experiments ^39^. The thermal peaks of both proteins indicate an exothermic transition with heat release, typical of protein unfolding transitions (*Figure 3, D*). The T_m_ values calculated in DSC are 58°C and 45°C for WT and I603M respectively, matching the values obtained in CD. The calorimetric enthalpy (ΔH_cal_) was calculated by integration of the area under the thermogram peak and resulted in 364 kJ/mol for C4 WT and 196 kJ/mol for C4 I603M. ΔH_v_ was determined by a fit of the thermogram to a two-state model of unfolding. Again, the ΔH_v_ (WT) is higher than ΔH_v_ (I603M), 330 kJ/mol and 222 kJ/mol, respectively (Table 5). The ΔH_cal_/ΔH_v_ ratio is 1.1 for WT and 0.89 for mutant, suggesting that the two-state model of unfolding is a reasonable approximation to the unfolding process of both proteins. The Gibbs free energy change (ΔG) of unfolding was calculated at 25°C considering ΔC_p_=0. The ΔΔG associated with the I603M mutation is 4.5 kcal/mol, confirming its highly destabilizing properties.

## 4. DISCUSSION

In the era of Next Generation Sequencing, an ever-increasing number of variants whose pathogenicity remains unclear is detected during genetic testing. In particular, many VUS that have no clinical value are identified ^16,18^. Indeed, the majority of genetic variants across all actionable genes are currently classified as VUS. This issue has become particularly challenging in the genetic diagnosis of HCM, one of the most common cardiac inherited diseases ^4,11^. Pathogenic mutations in HCM-associated genes are found in more than 60% of HCM patients, allowing cascade predictive genetic testing to early diagnose relatives who might be at risk of disease-related complications, such as sudden cardiac death. A different scenario follows identification of a VUS in the proband. Due to the absence of pathogenicity assignment of VUS, relatives cannot be reassured about their potential to develop HCM. Indeed, the Association for Clinical Genetic Science does not recommend predictive testing on family members following the finding of a VUS in an index case ^52^, which limits the utility of genetic testing in clinical practice. Functional assessment of VUS can help define their pathogenic nature. According to the ACMG, in the absence of enough genetic and population data, the most important criterion to establish causality of putative disease-causing mutations is a functional study to estimate the impact of mutations on a gene or protein function.

The c.1809T>G-p.I603M variant identified during genetic testing of an HCM patient was classified as VUS following ACMG criteria. Hence, the variant was selected for functional study to determine pathogenicity. We firstly evaluated if the c.1809T>G (p.I603M) variant induce alterations in RNA splicing. During splicing, introns are removed from primary transcripts, in a process that links together the flanking exons to generate mRNA ^47^. During the editing of pre-mRNA transcripts, the splicing machinery recognizes consensus sequences that also include the exons, particularly in the regions close to the intron-exon boundaries ^53^. Results show that c.1809T>G-p.I603M does not induce alterations in RNA splicing (*Figure 1 B, C*). Regarding protein stability, the variant does not perturb much the structure of the domain at 25°C, according to the far-UV and near-UV CD spectra (*Figure 3 A,B*). However, the thermodynamic stability of the C4 I603M mutant is severely compromised, as shown by lower T_m_ in thermal denaturation experiments by CD and DSC (*Figure 3 C,D*). Therefore, the p.I603M variant alters the stability of the central C4 domain of MYBPC3, leading to more frequent protein unfolding that can result in degradation of cMyBPC containing the mutation I603M ^54^. This scenario may lead to cMyBPC haploinsufficiency, which is believed to be a major pathogenic mechanism in truncating variants ^55^. Adding the functional analysis supportive of a damaging effect on protein to the ACMG criteria considered for this mutation, the pathogenicity assessment changes to likely pathogenic.

Remarkably, the variant p.I603M was identified in clinically affected subjects in combination with the p.T33RfsX15 truncating variant of cMyBPC (*Figure 1, A*). The two young subjects that have only the p.I603M do not show the phenotype although, according to our results, we do not exclude that they can develop the disease in the future. There is only one subject of the family that has cardiac hypertrophy in the absence of the mutation p.I603M, although the clinical manifestations probably are secondary to the coarctation of the aorta caused by an accident. In addition, there is one patient that carries only the truncated mutation and does not show HCM phenotype. This family reflects the difficulties associated with genetic testing in a clinical setting. In principle, frameshift mutations in MYBPC3 are thought to be causative of HCM ^27^. Hence, if p.I603M is also pathogenic, two of the members of the family would be compound heterozygotes for pathogenic mutations. This situation usually results in pediatric-onset HCM, which we did not observe in our case. This observation suggests that at least one of the variants is not highly damaging. Regarding p.T33RfsX15, there has been no study on whether N-terminal truncations in cMyBPC are as pathogenic as more central ones. The existence of re-start of translation downstream to the mutation-driven stop codon has been observed ^56,57^, which may lead to less damaging molecular phenotypes in these N-terminal truncations. Regarding p.I603M, it remains to be determined how the clinical severity of a mutation scales with the degree of destabilization, for which many pathogenic mutations need to be analyzed.

In summary, our study shows the advantages offered by functional assessment of mutations in the assignment of pathogenicity in the context of HCM. To fulfil its potential, we identify the need of extensive profiling of pathogenic mutations to find molecular phenotypes, such as protein destabilization ^46,58^, that can be used to guide functional assessment, a current research area in our groups.

## Acknowledgements

J.A.C. acknowledges funding from the Ministerio de Ciencia, Innovación y Universidades (MCNU) through grants BIO2017-83640-P (AEI/FEDER, UE) and RYC-2014-16604, the European Research Area Network on Cardiovascular Diseases (ERA-CVD/ISCIII, MINOTAUR, AC16/00045), the *Comunidad de Madrid* (P2018/NMT-4443), and the CNIC-Severo Ochoa intramural grant program (03-2016 IGP). The CNIC is supported by the Instituto de Salud Carlos III (ISCIII), MCNU and the Pro CNIC Foundation, and is a Severo Ochoa Center of Excellence (SEV-2015-0505).).

GF acknowledges funding from the Ministero dell’Istruzione, dell’Università e della Ricerca-RomePS35-126/IND.

