## Supplemental Figure 1 for "Thermodynamic destabilization informs pathogenicity assessment of a variant of uncertain significance in cardiac myosin binding protein C"

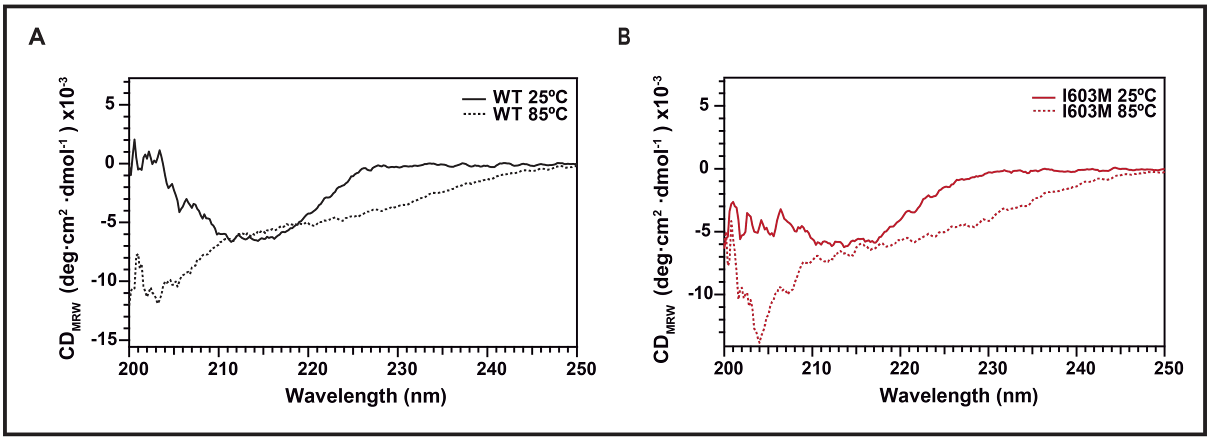


Supplementary figure 1: **Circular dichroism spectra of cMyBPC C4 WT and I603M before and after thermal unfolding.** CD spectra of C4 WT (A) and C4 I603M (B) monitored in the far-UV at 25°C (*solid line*) and 85°C (*dashed line*). Change in the shape of the far-UV spectra was observed when the sample temperature was increased to 85°C, indicating the unfolding of protein and loss of secondary structure for both WT and I603M. The maximum differences between spectra at 25°C and 85°C were observed at 230 nm. This wavelength was chosen for tracking the thermal denaturation of the proteins.
