## Supplemental Figure 2 for "Thermodynamic destabilization informs pathogenicity assessment of a variant of uncertain significance in cardiac myosin binding protein C"

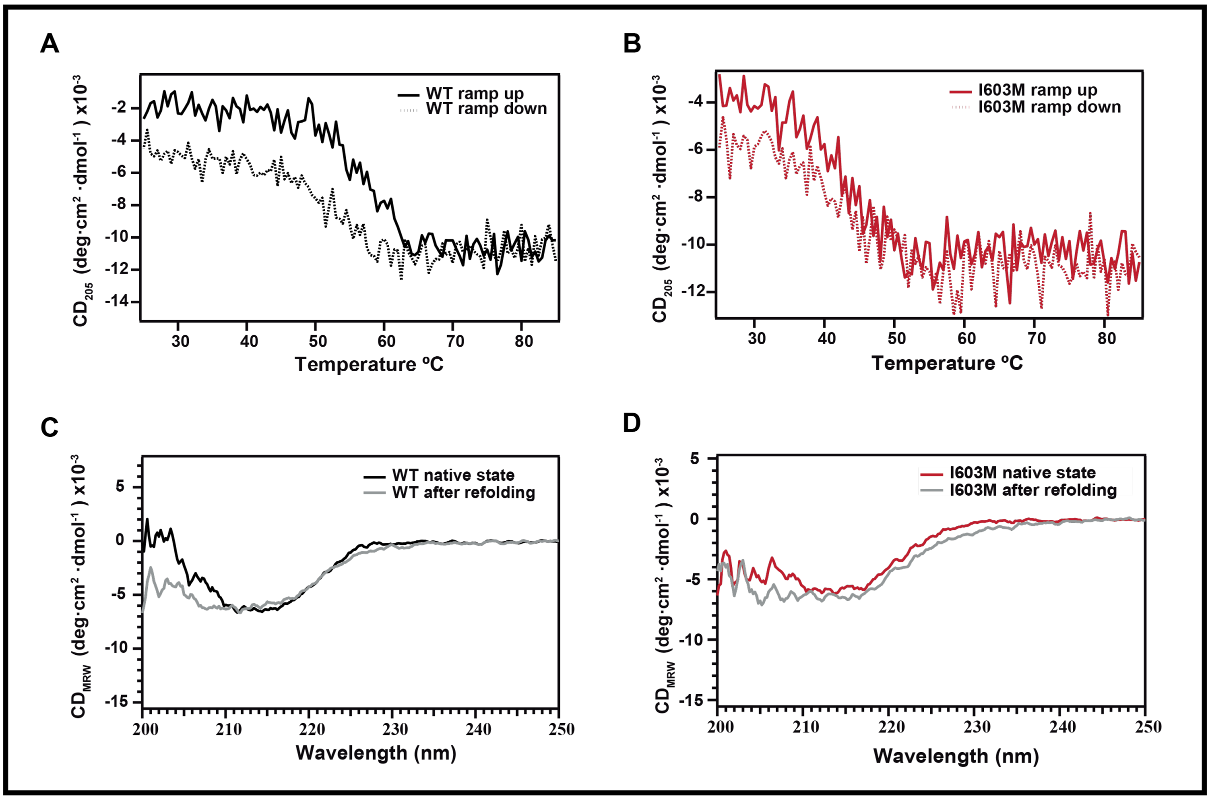


Supplementary figure 2: **Characterization of the reversibility of the thermal denaturation of the C4 domain.** Thermal denaturation (*solid line*) and renaturation (*dashed line*) of C4 WT (A) and C4 I603M (B) collected as change of ellipticity during thermal ramp up (from 25°C to 85°) and thermal ramp down (from 85°C to 25°C). The far-UV CD spectra of C4 WT (C) and C4 I603M (D) of protein before and after (*grey traces*) thermal denaturation/renaturation protocols.
